# Deformability Cytometry Clustering with Variational Autoencoders

**DOI:** 10.1101/2022.10.01.510460

**Authors:** Daniel D. Seith, Cody Combs, Zuzanna S. Siwy

## Abstract

Mechanobiology has shown great success in revealing complex cellular dynamics in various pathologies and physiologies. Most methods for assessing a cell’s mechanical properties, however, generally extract only a few physical constants such as Young’s modulus. This can limit the potential for accurate classification given the wide variety of rheological properties of cells, there are many ways for cells to differ. While it was recently shown that deep learning can classify cells more accurately than traditional approaches, it is not clear how this may be extended to unsupervised classification. In this work, we showcase the potential for a deep learning model to classify cells in an unsupervised fashion using a blend of physical properties. We introduce the combination of a variational autoencoder and a previously described clustering loss for classifying cells in an unsupervised fashion.

## Introduction

The ability to characterize a cell’s mechanical properties is emerging as a new method for high throughput characterization. One of the most commonly used mechanotyping method is deformability cytometry and has been shown to characterize COVID-19 pathology,^1^ red blood cell disorders,^2^ and circulating tumor cells.^3^ While there are previous works applying deep learning to deformability cytometry,^4,5^ there are few, if any, instances of the application of unsupervised deep learning to deformability cytometry. Given deep learning has shown to be particularly suited to vision tasks, we wondered if it would also unlock new capabilities for deformability cytometry. In this work, deep learning models for unsupervised deformability cytometry classification are investigated. The dataset for this study was previously described and includes HL60 cells before and after treatment with Cytochalasin D (CytoD).^6^ CytoD is known to disrupt actin polarization and leads to an increased deformability of HL60 cells.^7^ Unsupervised cell classification could be investigated using two routes (Figure 1). In the first approach, scalar features, such as region deformation, relaxation rate, and peak deformation, can be extracted manually or with a feature extraction algorithm, and subsequently utilized by a clustering algorithm. In a second approach, deep learning methods can be used to better access the high dimensional latent space for clustering.

**Figure 1:**
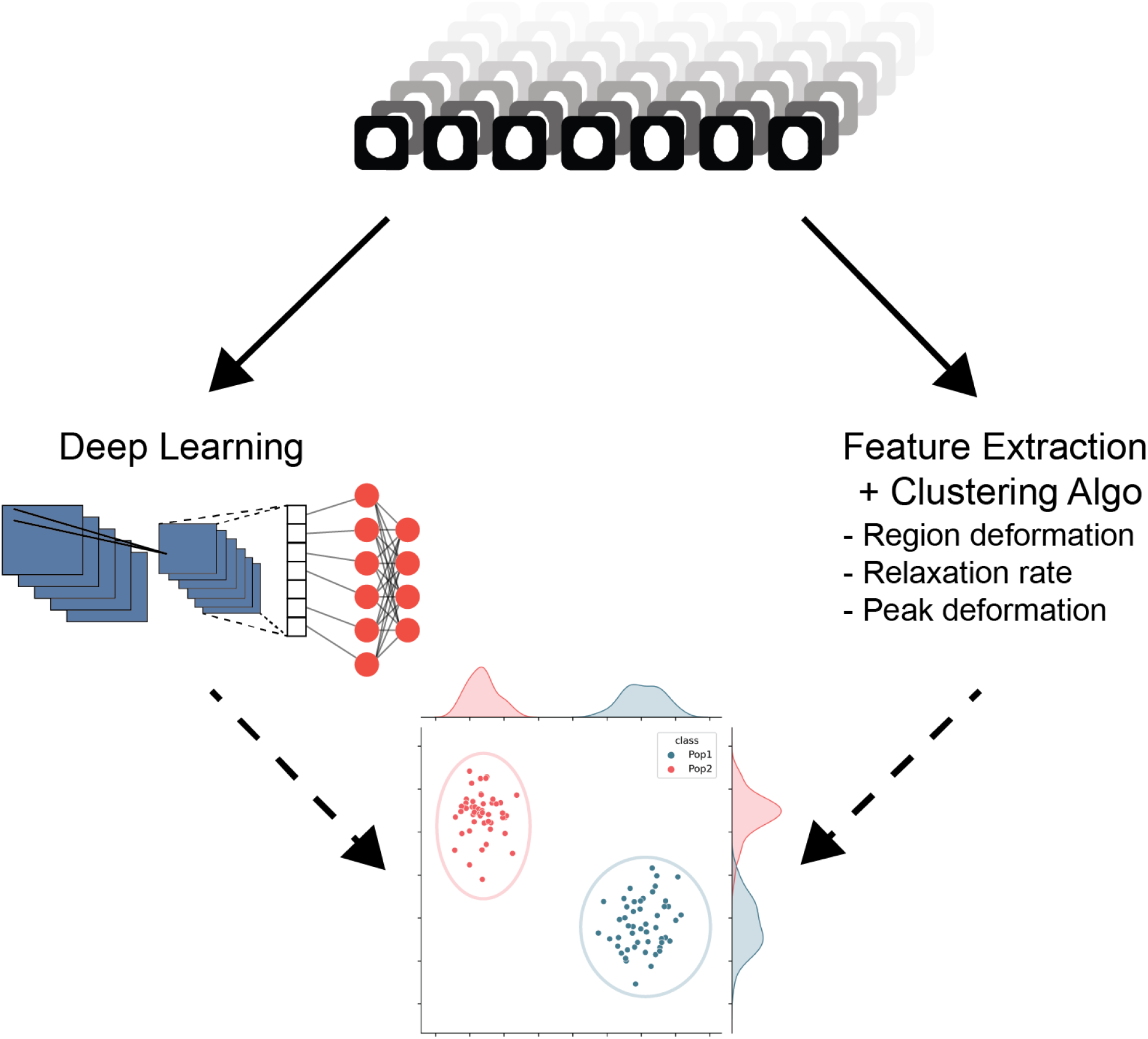
Routes to clustering DC data. Two potential routes to clustering deformability cytometry data exist and are shown here. Each provides unique strengths and weaknesses.

Unsupervised clustering is first investigated using traditional clustering algorithms: a Gaussian mixture model (GMM) and spectral clustering algorithm, with principle component analysis (PCA) extracted features. However, these scalar features contained a very significant class overlap and the clustering task was too challenging for the clustering algorithms to accurately separate the two populations of cells, HL60 and HL60d. These issues motivated us to look into deep learning models for clustering, namely a variational autoencoder (VAE), which has previously shown success in clustering cell data.^8,9^

Here, a novel unsupervised clustering algorithm using a VAE for deformability cytometry was developed, which is referred to as DeepDeform. A VAE alone was tested first as a benchmark of clustering performance. However, the VAE was limited in its ability to cluster cell populations, since it is only capable of compressing data and is not inherently a classifier. We then assessed the performance of the VAE in tandem with the previously mentioned traditional clustering algorithms, which showed poor clustering performance likely due to improperly balanced clusters.^10^ In DeepDeform, a semantic clustering by adopting nearest neighbors (SCAN) loss is integrated to abrogate the deficiencies of traditional VAEs. The implementation of SCAN enabled better access to the high dimensional latent space for clustering cell populations, since it did not require compression of the latent space. Using DeepDeform, the clustering accuracy of the two cell populations rivaled that of several supervised methods.^11^ Additionally, the accuracy approached that of supervised models when selecting confident samples.

The manuscript is organized as follows. Section 0.1 provides an overview of the methods, such as the dataset and training procedures. Section 0.2 describes initial efforts to investigate cell populations using conventional clustering and dimensionality reduction techniques, such as principal component analysis (PCA). Section 0.3 presents the development, optimization, and performance of unsupervised clustering using DeepDeform on the HL60 model system.

## 0.1 Methods

### Channel Design

The dataset for unsupervised clustering was obtained using the same microfluidic channel design as described previously.^6^ Briefly, the channel was composed of four zones: two narrows each followed by two cavities. Specifically the channel was 150 μm long with equal length sub-regions (50 μm). The narrow regions have widths of 25 μm. The first cavity has a width of 50 μm and the second cavity is the bulk. The height of the channel was 20 μm.

### Dataset

The dataset for unsupervised clustering included 2239 human leukemia 60 (HL60) cells. The two classes are comprised of untreated (HL60) and cytochalasin D (CytoD) treated HL60 cells (HL60d). There is a roughly even split (51% - 49%) between the two classes: HL60 and HL60d. CytoD, a fungal toxin, disrupts actin polymerization, resulting in a decrease in cell stiffness, while leaving morphology and radius relatively unaffected.^6^ This means that any clustering algorithm needs to rely heavily on mechanical properties to separate the populations rather than more accessible information such as a cell’s morphology or radius. Since the class labels are known, an unsupervised model’s ability to extract these known labels can be evaluated without the model training on them directly. There are 4 different biological replicates of HL60 cells and 4 biological replicates of HL60d cells. Multiple technical replicates were also gathered with the same density of cells being used for each session. The data gathered consist of spatially synchronized 35 frame sequences of 96 × 96 binary masks of cells passing through the channel. The video data can be represented in fewer dimensions by identifying important features such as peak deformation, change in deformation in the first and second regions, as described previously.^6^

### Features

The features for the unsupervised clustering were extracted from the raw binary masks.

### VAE Architecture

Provided below is the VAE architecture of DeepDeform. Raw binary mask data were used from the previous work. Adam was used as an optimizer with a learning rate of 1e-3. Gradients were clipped at a norm of 3.0. Experiments were tracked using wandb.^12^ The loss function was the sum of reconstruction loss and KL divergence. Both training and validation losses were monitored during training in addition to the average class of the five nearest neighbors (Figure 2). The batch sizes were 32. Data were shuffled each iteration. Unless specified, all dense and convolutional activation functions were leaky rectified linear activation unit.

**Figure 2:**
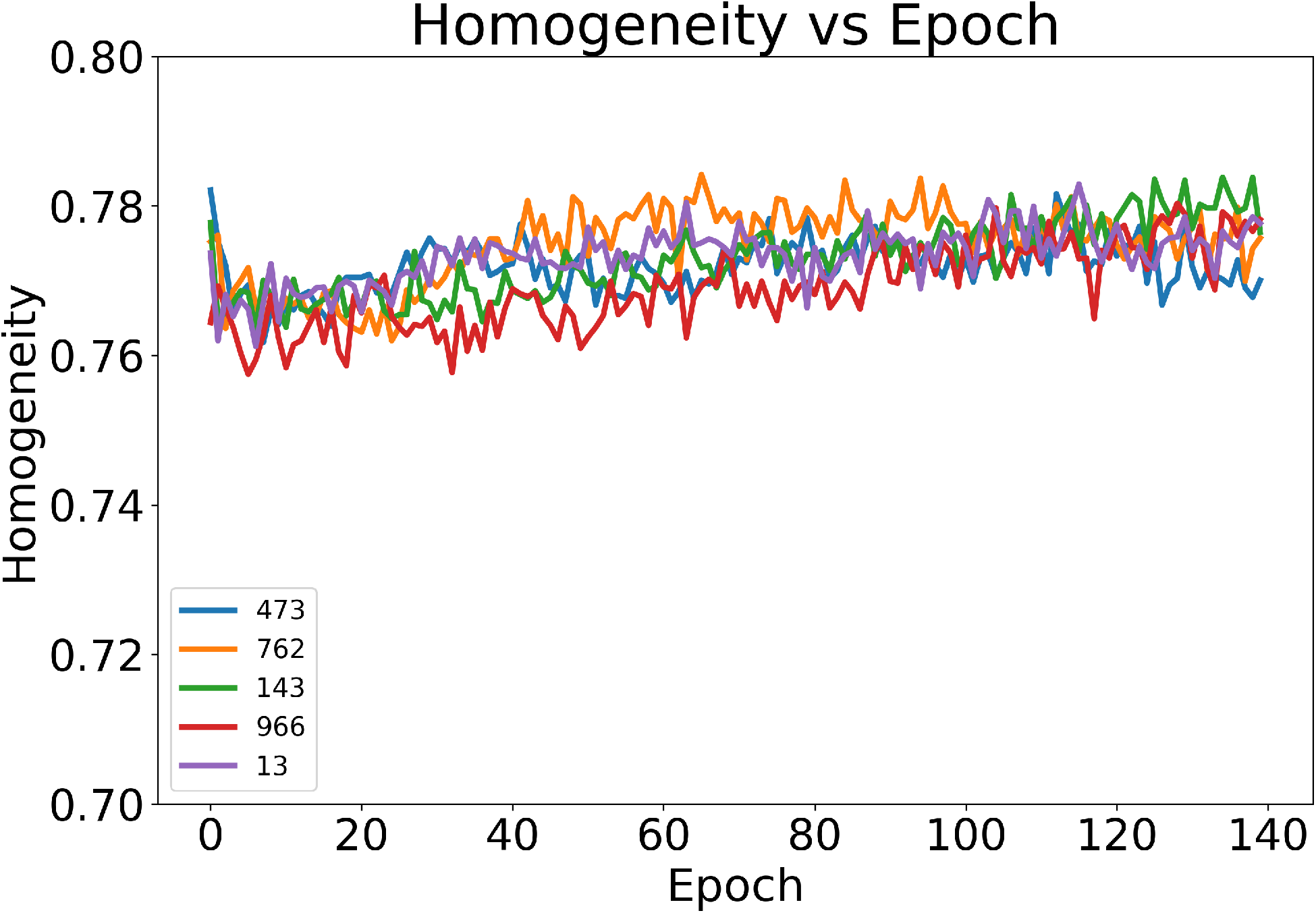
Average homogeneity of all points’ nearest 5 neighbors during VAE training.

### Encoder Network

1. Conv2D - Filters: 8 - Kernel: 3×3
2. MaxPooling2D
3. Conv2D - Filters: 16 - Kernel: 3×3
4. MaxPooling2D
5. Conv2D - Filters: 32 - Kernel: 3×3
6. MaxPooling2D
7. Flatten
8. GRU - Dimension 32

### *μ* Network

1. Dense - Dimension 8

### *σ* Network

1. Dense - Dimension 8

### Decoder Network

1. Repeat Vector - Number of repeats: Number of input timesteps
2. GRU - Dimension 32
3. Dense - Dimension 24 × 24 × 64
4. Reshape - 24 × 24 × 64
5. Conv2D Transpose - Filters: 64 - Kernel: 3×3
6. Conv2D Transpose - Filters: 32 - Kernel: 3×3
7. Conv2D Transpose - Filters: 16 - Kernel: 3×3

### Semantic Clustering by Adopting Nearest Neighbors (SCAN) Implementation

Once encoder training was completed, the model weights were frozen and a single two dimensional, softmax-activated, dense projection layer was added. The SCAN loss was implemented according to the work of Gansbeke et al.^13^ with two separate Adam optimizers for the consistency and entropy components respectively. Both optimizers had a beta_1_ of 0.9, a beta_2_ of 0.999 and an epsilon of 1e-7. The learning rate of the consistency optimizer was set to 1e-3 while the entropy optimizer was 5e-3.

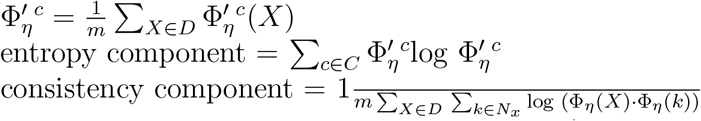

### Code Availability

The python scripts implemented for data processing and training machine learning models are available at: https://github.com/siwylab/vae_dc

## 0.2 Unsupervised Clustering Using Manually Extracted Features

In our previous work we observed that for a supervised task, machine learning was required to extract the most useful information out of cell cytometry data.^6^ Given that many problems in science are open-ended and do not have known labels, it is necessary to extend our deformability platform to unsupervised problems. It is not, however, clear what the ideal route towards creating a clustering algorithm is (Figure 1). One route, utilizing manually extracted features, could offer a simple and interpretable model for classification. Meanwhile, deep learning utilizes the raw binary masks and could offer improved clustering accuracy.

To investigate the first route, the ten manually extracted features (Table 1) previously found useful for supervised classification were first considered. These data were a good starting point for the analysis since they have a much lower dimensionality than the sequence of raw masks (10 vs. 35 × 96 × 96). However, we condensed the data further since most clustering algorithms show best performance with two or three dimensions. The dimensionality was further reduced down to two dimensions with PCA, which is a lightweight and accessible technique for dimensionality reduction. With a two-dimensional input space, the practicality of existing clustering algorithms: Gaussian mixture model (GMM) or the spectral clustering model was assessed with the data. Both algorithms are easily accessible and have been previously used for clustering biological data.^14,15^ The desired task was to learn patterns inherent to the data that would allow them to correctly assign class labels (HL60/HL60d) without training with the labels directly. The compressed features are shown in Figure 3. Both GMM and spectral clustering showed poor performance with a clustering accuracy of only 55.4% and 50.0%, respectively. This poor performance was due to the large overlap between the two classes in the lower dimensional space created by PCA. Additionally, more class overlap likely existed in the hand-picked features than in the binary masks.

**Table 1:**
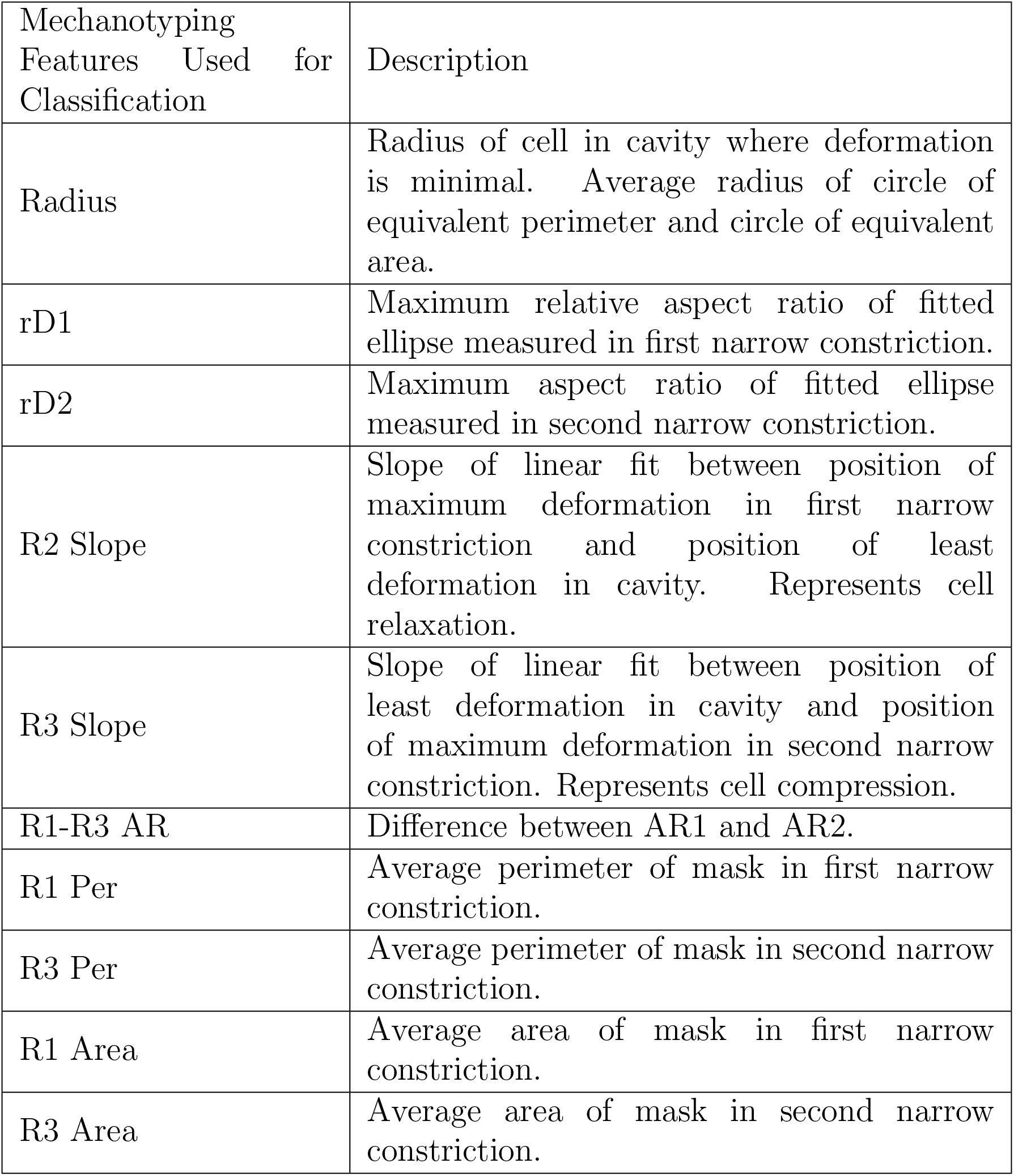
Parameter table

**Figure 3:**
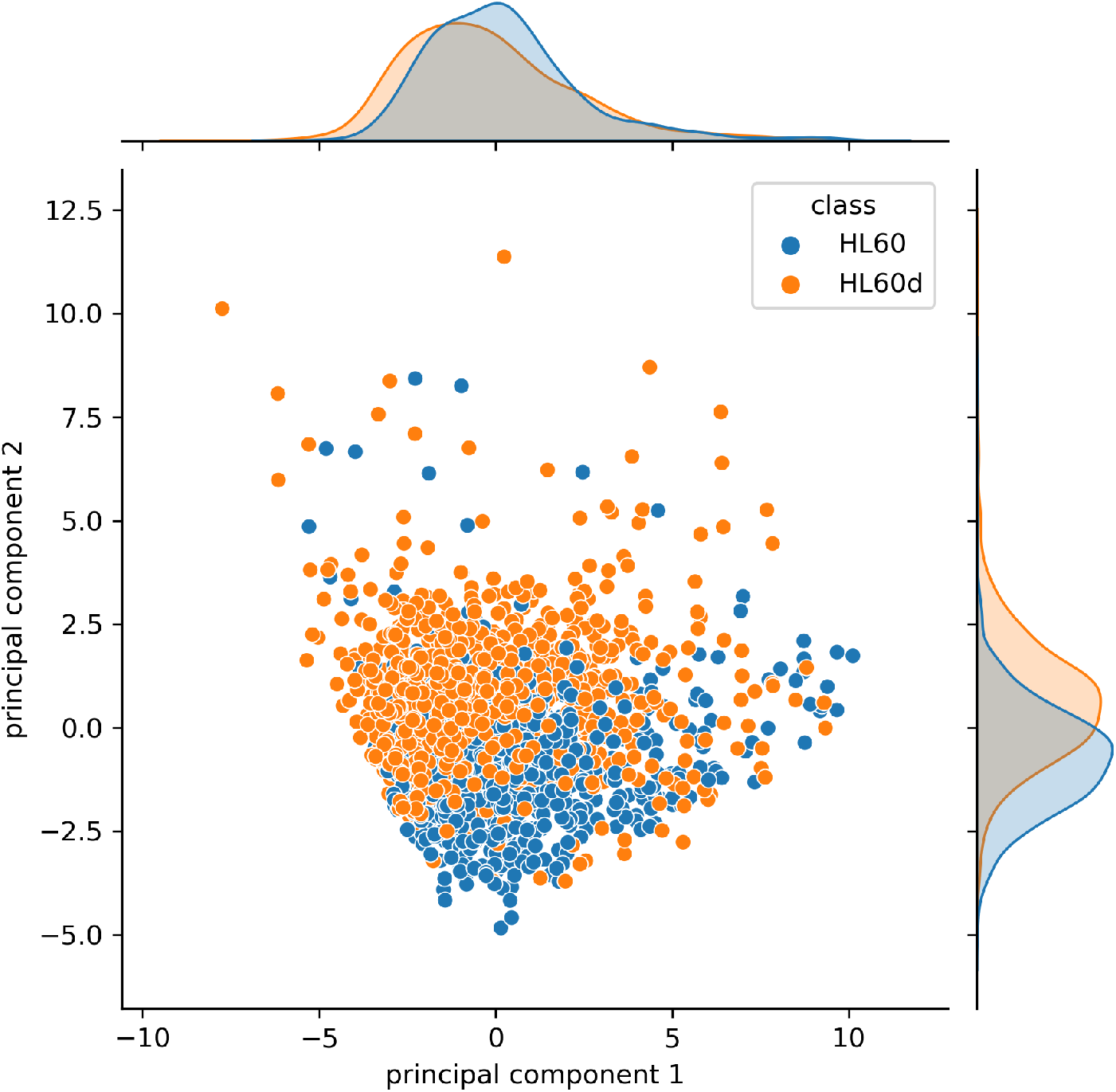
First two principal axes from PCA, ground truth labels are shown.

## 0.3 Unsupervised Clustering Using Deep Learning

It was hypothesized that an optimized dimensionality reduction technique would compress the input data more effectively than manually extracting features. This would enable the use of richer data, such as the raw binary masks. A variational autoencoder (VAE) was therefore applied as an algorithm to access the rich latent space created by efficient compression of binary masks. VAEs reduces the dimensionality of input data to arbitrary dimensions^16^ and has been previously used for clustering cell data.^8,9^

### VAE with Traditional Clustering

To maximize the potential of the model, the raw sequence of binary masks are used as inputs, which contained more data than the manually extracted features. The VAE was created using a convolutional recurrent encoder and a corresponding decoder. The decoder network contained a recurrent layer and used convolutional transpose layers. This allowed us to scale up the decoder input data and create an output size equal to the raw binary masks. The full VAE architecture is provided in Figure 4. The VAE was trained on the sum of the reconstruction loss and the Kullback-Leibler divergence (Equation 1) with an encoder output size of 32 dimensions. This size was chosen through a hyperparameter sweep that optimized the loss of the VAE.

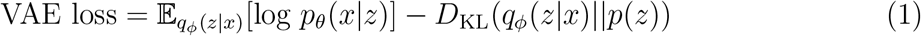

**Figure 4:**
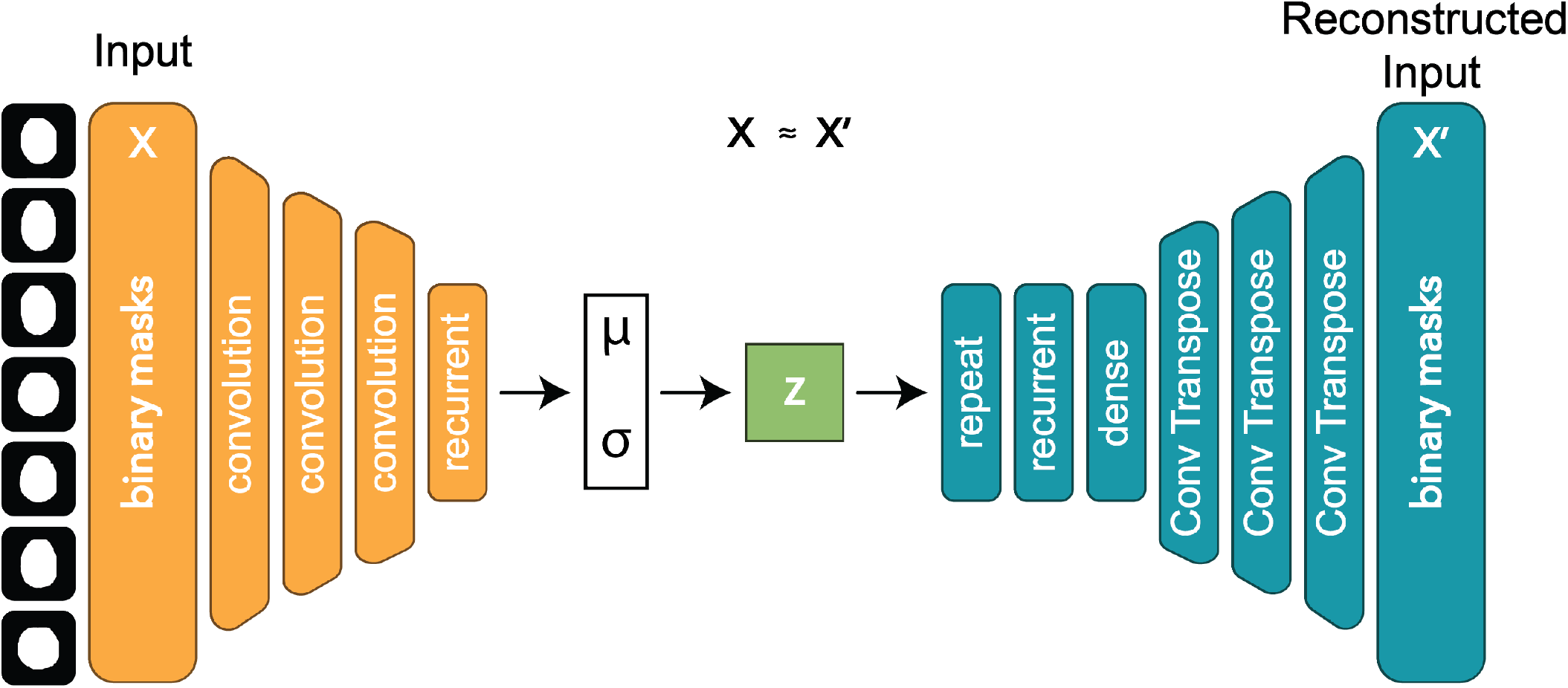
Application of VAE architecture for reconstructing the sequence of binary masks. The encoder network is colored orange, and the decoder network is colored blue.

The GMM and spectral clustering models were implemented in order to utilize this latent space created by the encoder. The results are provided in Figure 5. Since the latent space created by the encoder was still too high-dimensional (32-dimensional) for the clustering algorithms to be used on their own, PCA was subsequently applied to further reduce the dimensionality to two. Both clustering algorithms tested showed an improved mean accuracy but noisy performance (±3.5%). The unstable performance could be caused by an insufficient balance of class assignments.

**Figure 5:**
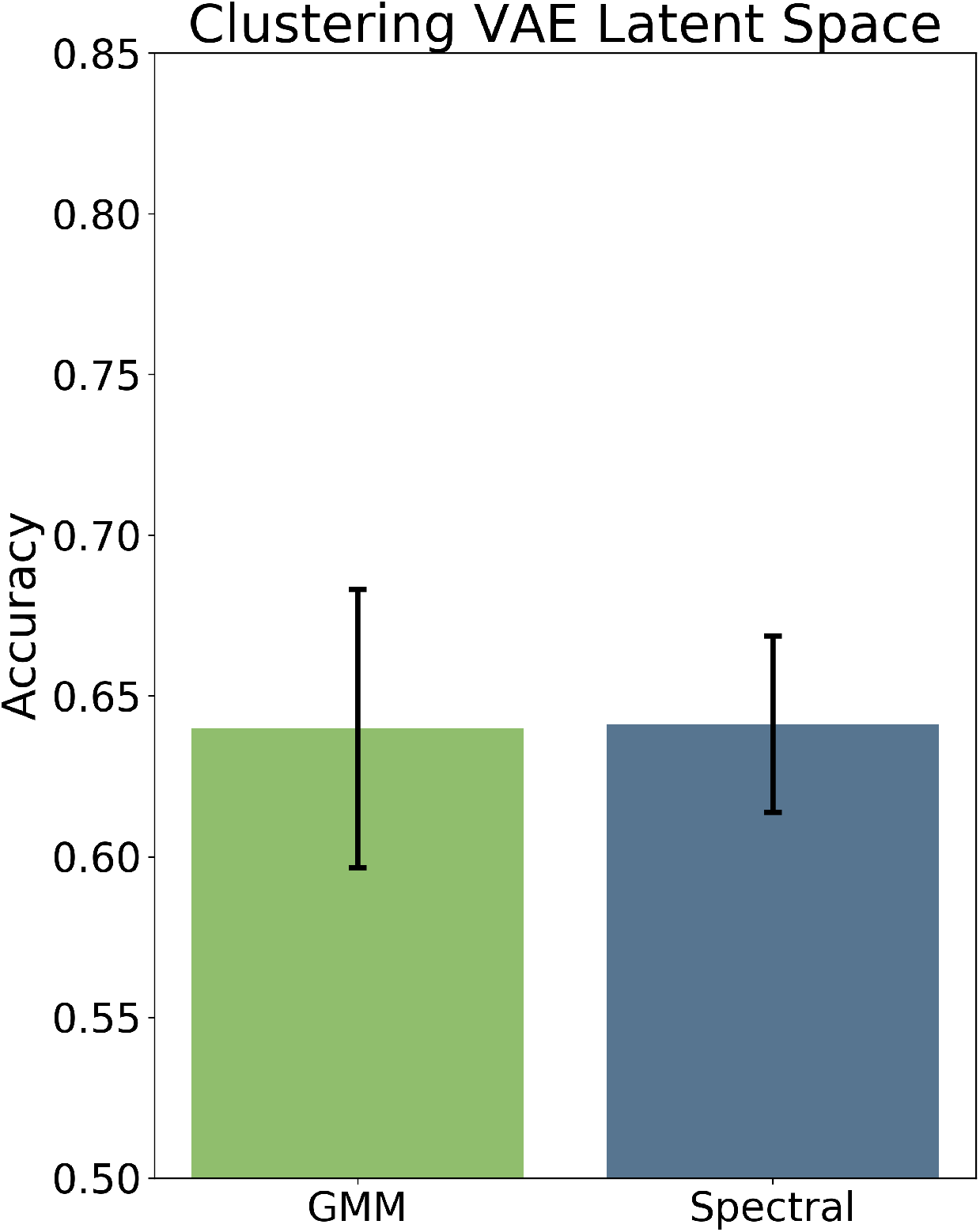
Accuracy of GMM and spectral clustering on trained VAE latent space. Five different random seeds are tested.

To seek improvements, the data were carefully studied. Interestingly, we noticed that each point contained a high nearest-neighbor homogeneity, meaning that a point was likely to be surrounded by other points belonging to the same class. We realized that such structure in the latent space could be harnessed to improve clustering accuracy. Specifically, a technique designed by Gansbeke et al.^13^ was shown to take advantage of such a structure, applied in the context of unsupervised image clustering. Their method, among other features, uses a loss function (Equation 1) that encourages balanced and confident class predictions. We hypothesized that this loss function could also allow the encoder to cluster our data with greater accuracy. The VAE model was then trained and the encoder weights were frozen. A projection layer was then appended to the encoder and the network was trained on the SCAN loss function (Figure 6). The balance between the two components of the SCAN loss (entropy and consistency) was then optimized.

**Figure 6:**
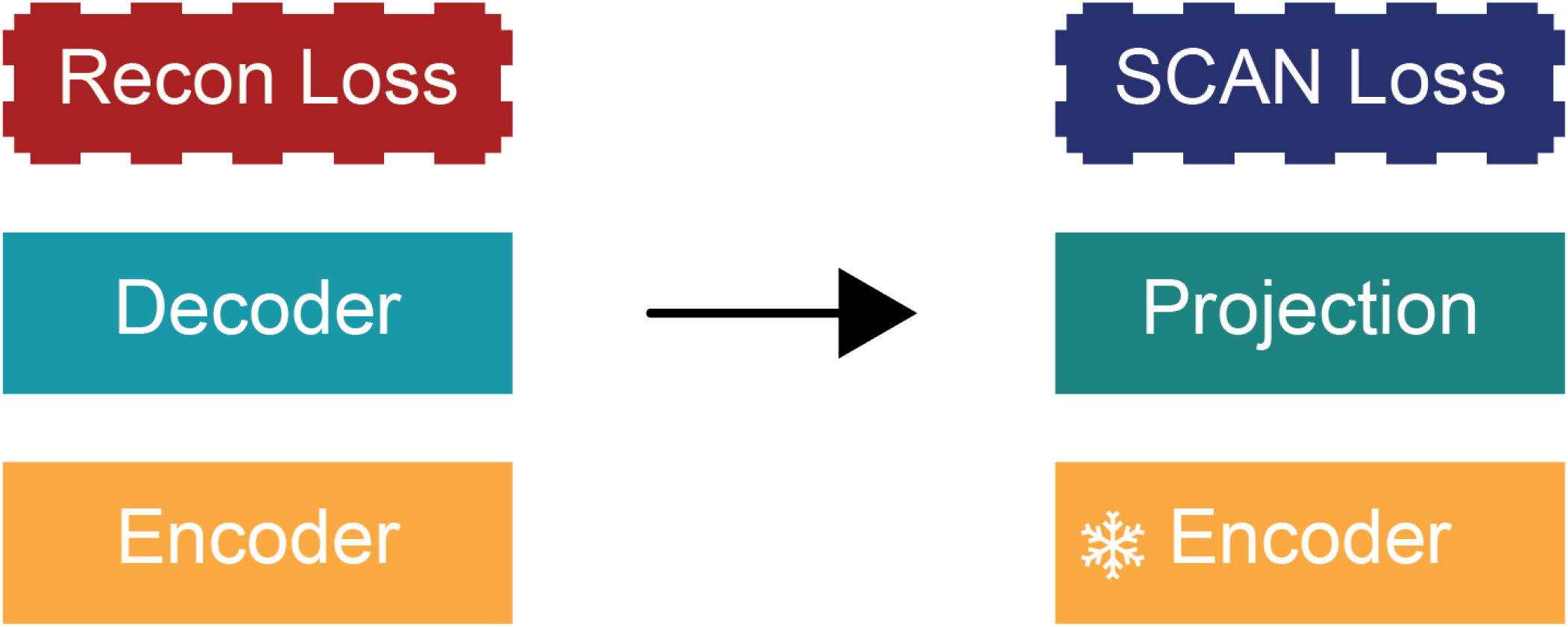
The scheme of the training procedure. The encoder model is first trained in conjunction with a decoder using standard VAE losses. The encoder weights are then frozen and a projection layer is appended. This projection layer is then trained using the SCAN loss.

### Initialization Sensitivity Tests

The next step was to characterize the performance of this trained clustering algorithm. In previous reports, deep unsupervised clustering techniques have been observed to be sensitive to initialization.^17,18^ Therefore, initialization was the first component of the performance to be tested. We tested the model’s sensitivity with five random seeds. The sweep over the five random initializations for the total training, VAE loss and then SCAN loss, resulted in an average accuracy of 61.9 ± 4.2%. As shown in Figure 7, the model was still quite sensitive to initialization.

**Figure 7:**
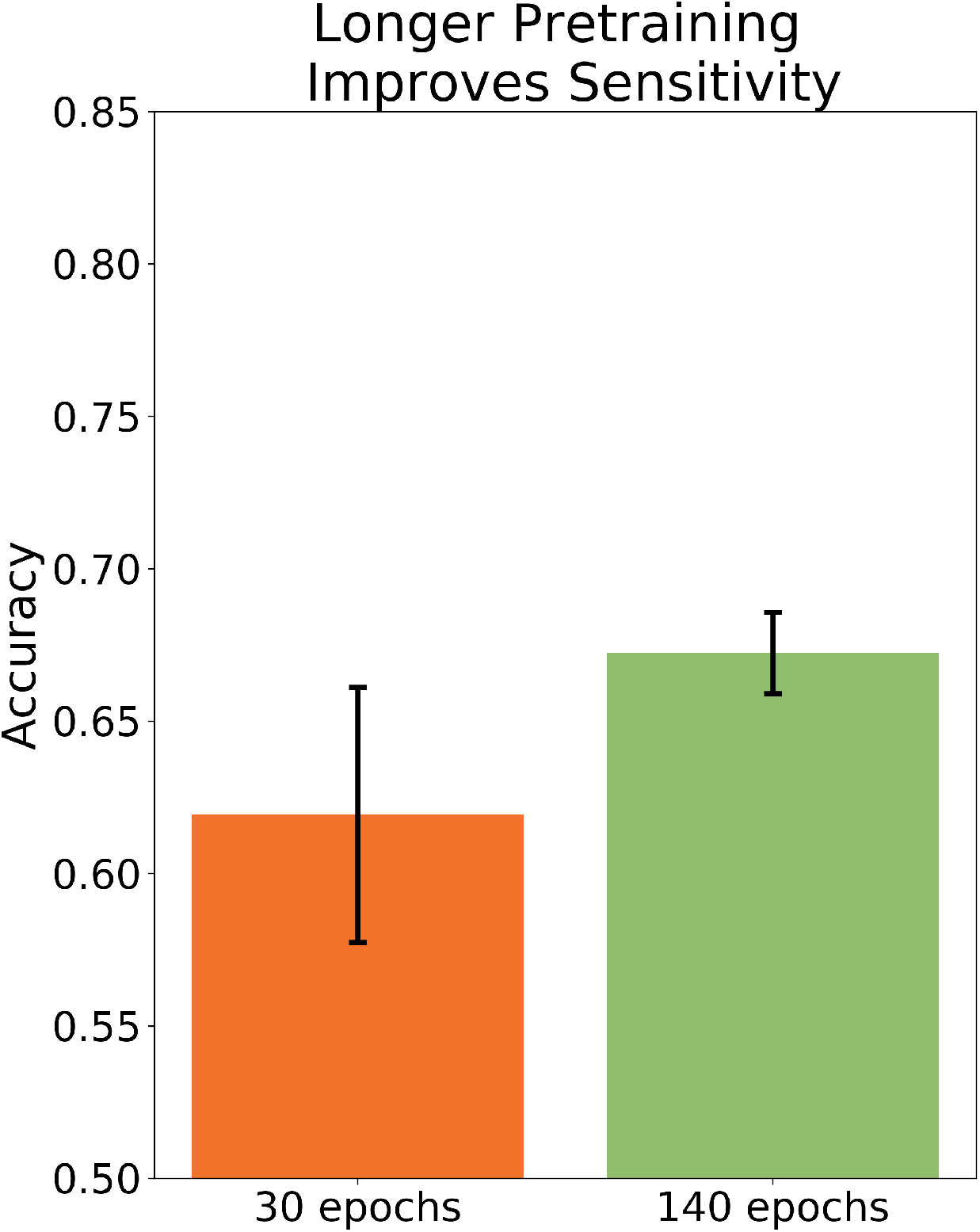
Longer training improves sensitivity. The accuracy of two training sweeps over 5 random seeds are averaged and shown.

We have also realized that the poor classification performance could be due to a global structure rather than a local one. Specifically, while the five nearest neighbors generally belonged the same class, the global structure may not be conducive to clustering and is not reflected in the average nearest neighbor homogeneity. We indeed observed that increasing the VAE training time from 30 to 140 epochs was very beneficial. This extended training decreased both the standard deviation in accuracy from 4.2% to 1.3% (Figure 7) and increased the mean accuracy from 61.9% to 67.2%.

To further mitigate noise and improve training stability, we probed a correlation between each of the losses (entropy, consistency loss) and clustering accuracy. If we observed that one of the losses (whose value does not require knowledge of the ground truth label) could be used as a proxy for clustering accuracy, multiple random seeds could be tested facilitating selection of a high-performing model, and an improved overall performance. To investigate a possible correlation, the encoder was trained with five different random seeds (Figure 8). Although no clear correlation between losses and accuracy was observed, it was hypothesized that clustering losses should be compared within a given encoder model, not necessarily between them (i.e. an individual model could be optimized through random seed sweeps for the second step, though not necessarily the first). Next, for each encoder model, three different random seeds were used to train on the SCAN loss. However, none of the loss functions substantially correlated with the accuracy of the clustering, which prevented the selection of a performant model (Figure 9).

**Figure 8:**
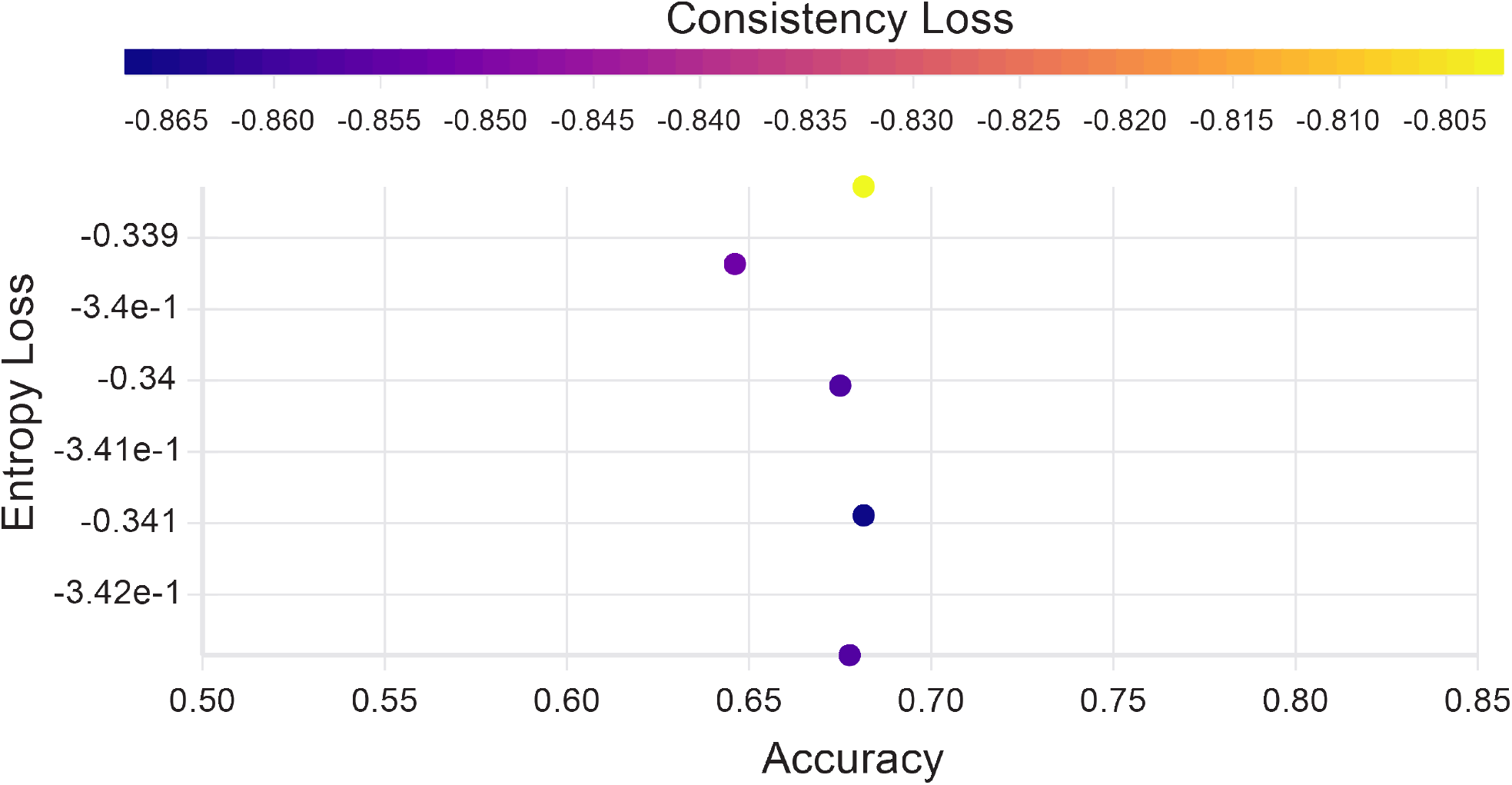
No relationship between SCAN losses and accuracy for a single frozen encoder. A random seed sweep was conducted using five random seeds. Classification accuracy is plotted against entropy and consistency losses.

**Figure 9:**
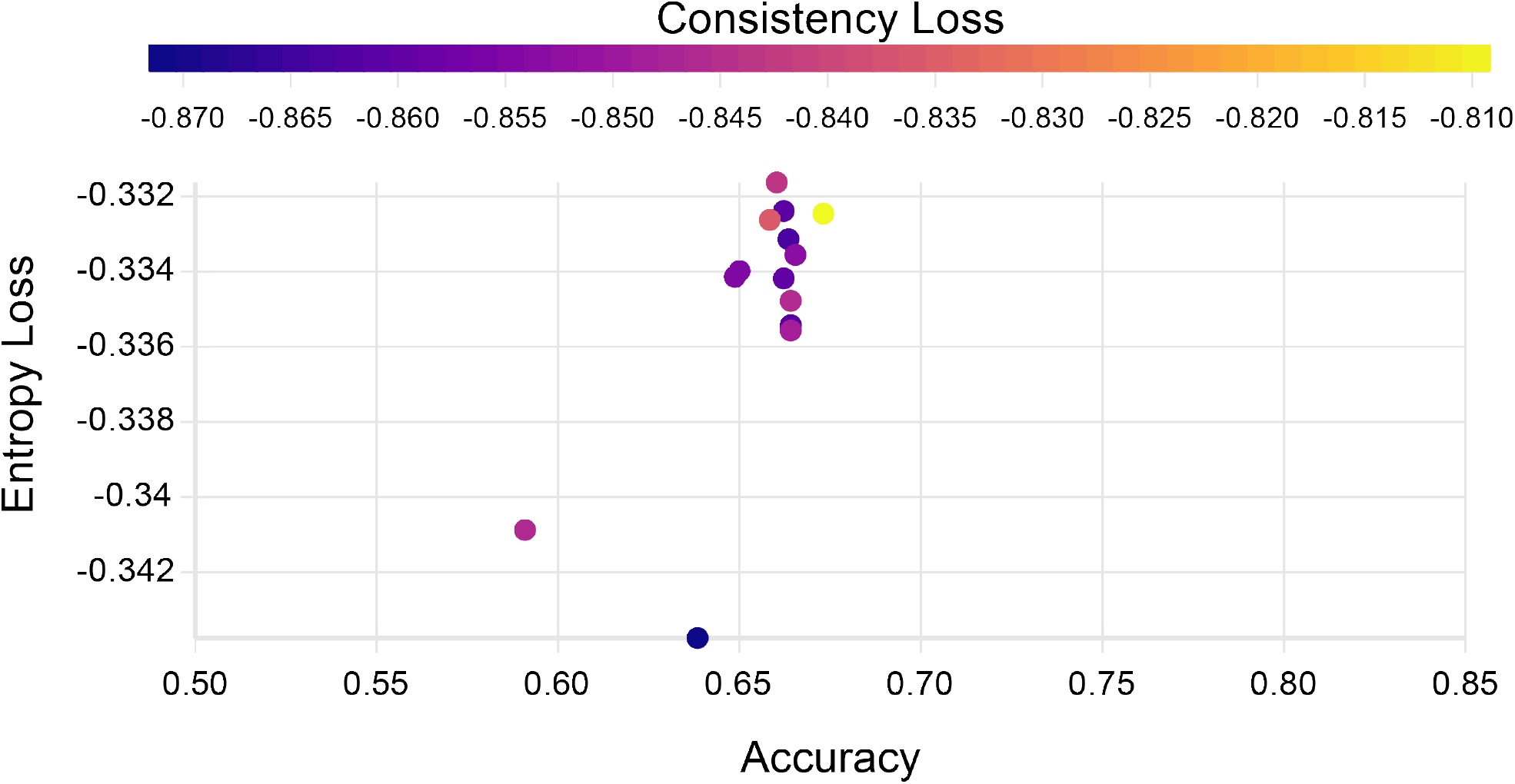
No relationship between SCAN losses for differently seeded encoders. Each of the encoder models from figure 8 is used to initialize the clustering model. For a single encoder, three different random seeds were used in an effort to find an optimal clustering.

### Augmentations and Selecting Confident Samples

In an effort to improve the model’s clustering accuracy and encourage the encoder to extract semantically relevant features, the strategy of augmenting data was considered. This strategy seeks to modify the data while leaving the underlying class of the data easily discernible, forcing the model to rely on higher level features. Since the choice of augmentations is generally task-dependent^19^ augmentations were chosen that destroyed low-level information while preserving high-level information, such as the overall order of timesteps. To test the potential of this approach, we used several augmentations during training (Figure 10). The four augmentations provided in Figure 10 include a randomly placed box of zeros of size 30 × 30 (style 1), complete subtraction of a given image (style 2), zeroing out 33% of the pixels (style 3), setting 33% of pixels to one (style 4), and shuffling of a given image with the next two images in the sequence (style 5). Samples were chosen at random to be augmented by any one of the listed styles. If a sample was chosen for augmentation, 10% of the timesteps were augmented with the randomly chosen augmentation style. It was observed that the augmentations actually decreased the performance significantly (Figure 0.3).

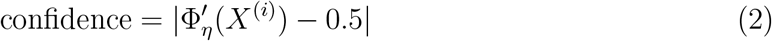

**Figure 10:**
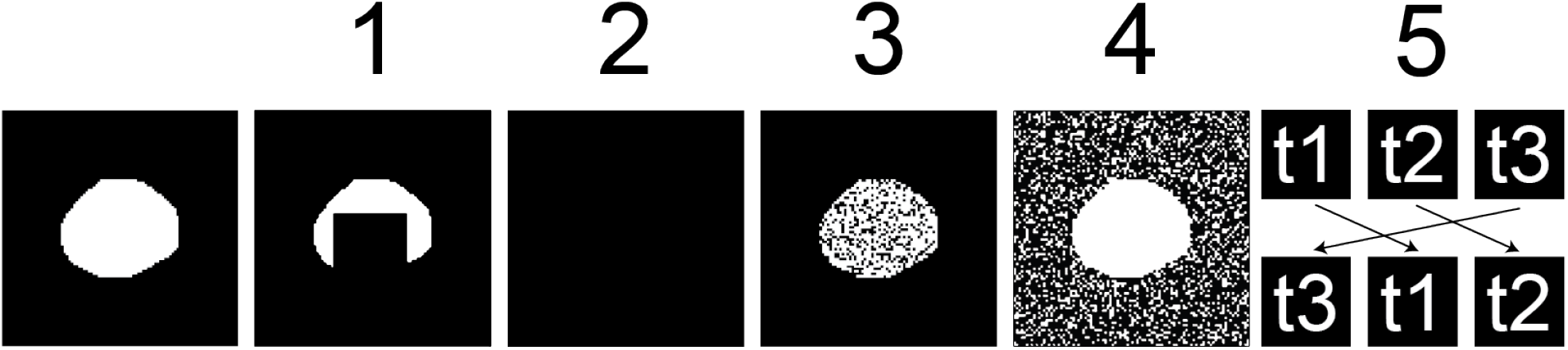
Five different augmentation styles. The first image on the left shows a sample image before augmentation. Style one shows a randomly placed box of zeros of size 30 × 30. Style two shows the complete subtraction of a given image. Style three shows 33% of the pixels zeroed out. Style four shows 33% of pixels set to one. Style five shows the shuffling of a given image with the next two images in the sequence.

**Figure 11:**
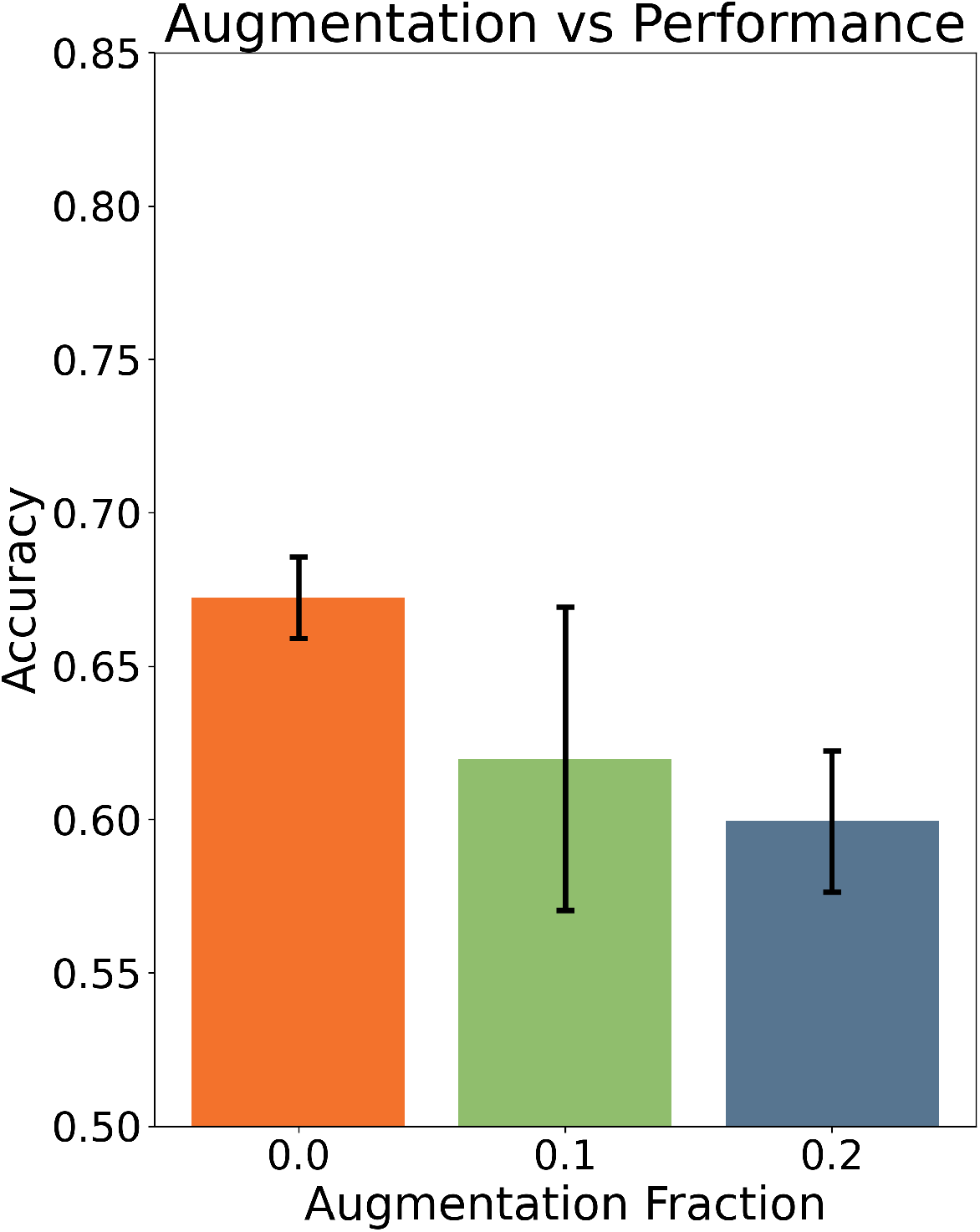
The fraction of augmented samples versus the classification accuracy. If a given sample is chosen to be augmented, on average 30% of the time steps will be augmented. Each of the five augmentations is selected at random for each timestep.

After inspecting model performance more closely, we observed that confidently clustered samples were often accurate (prediction confidence defined in equation 2). Although one generally needs to work with the whole dataset, it can be valuable to select representative examples with a high likelihood of being a true positive. For example, in enriching a cell population for downstream processing, precision is the most important metric. Therefore, we sought to identify examples to represent each cluster by selecting only confident samples (*>* 0.495). The top predictions made up ∼10% of all predictions with an even class distribution (Figure 12). Setting a threshold for classified samples improved the accuracy to ∼80% (Figure 13). For comparison, the accuracy of a supervised deep learning model trained on binary masks is 91% and the accuracy is 74.5% for a supervised SVM trained on hand-picked features.^6^

**Figure 12:**
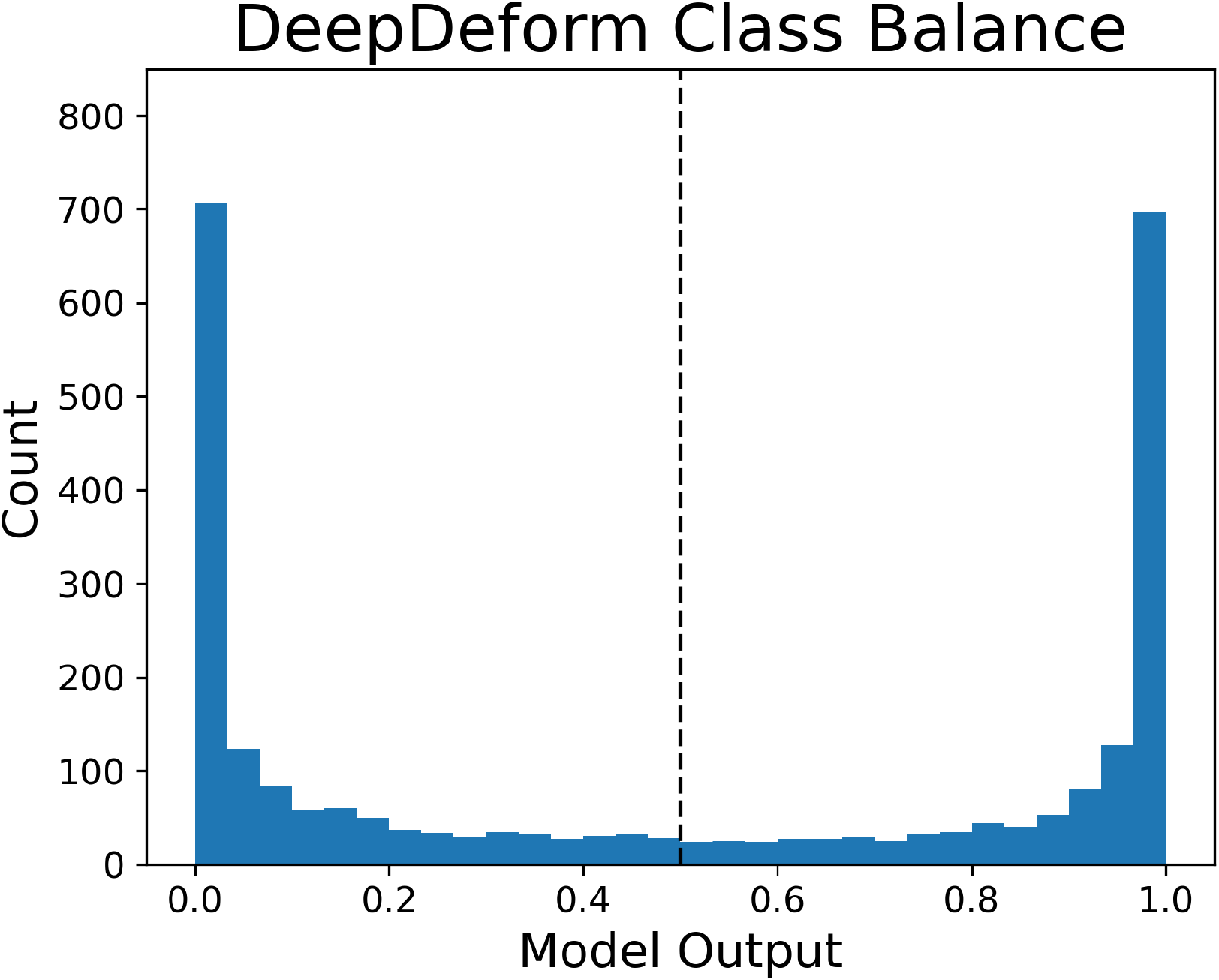
Averaging class balance over five random seeds. A model output of zero indicates a confident prediction of HL60 while a prediction of one indicates a confident prediction of HL60d. A prediction of 0.5 indicates complete uncertainty, which is illustrated with a dotted black line.

**Figure 13:**
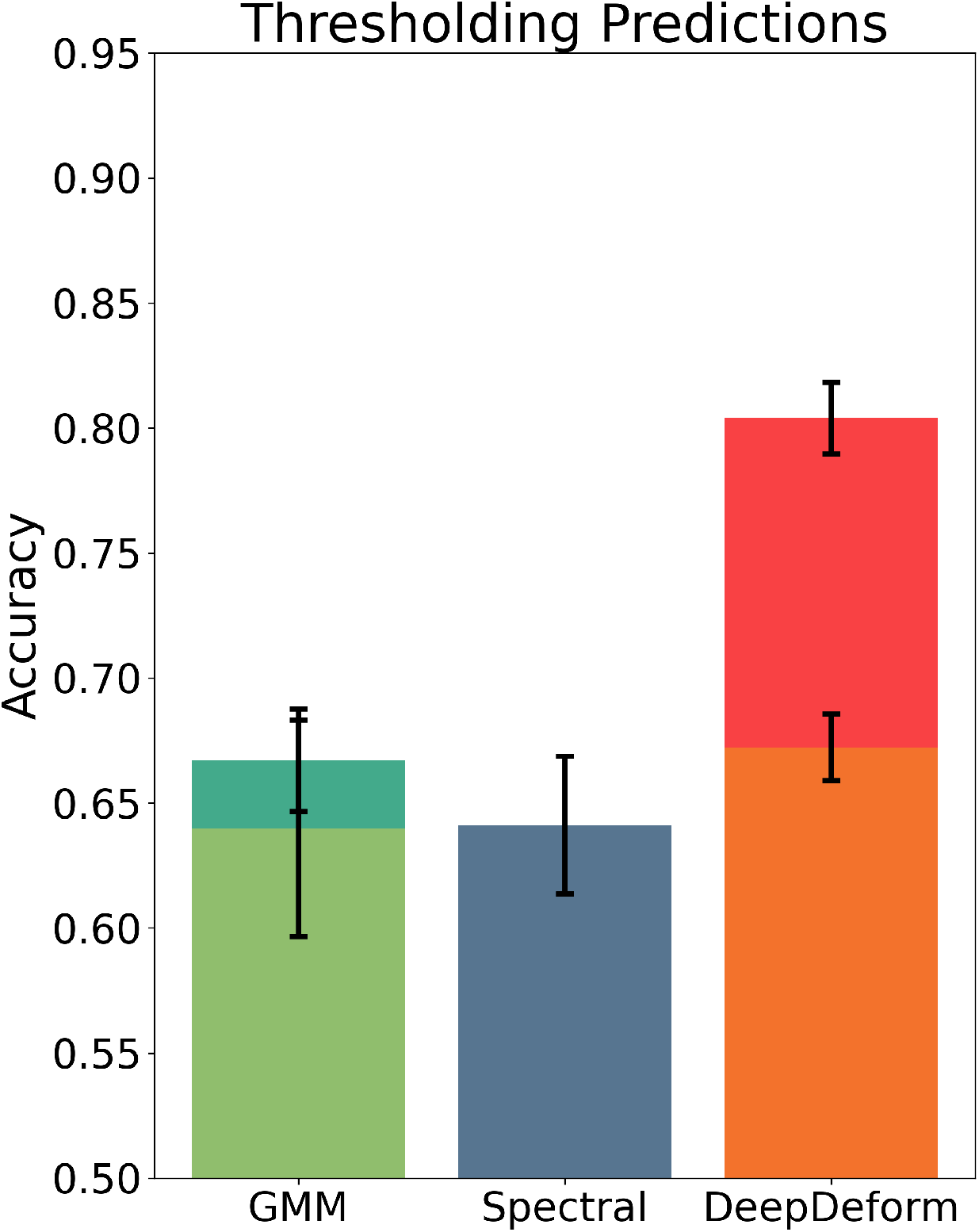
Selection of predictions with a confidence over 0.495 for both the GMM and DeepDeform. The dark green bar and red bars reflect the selection of confident samples for the GMM and DeepDeform, respectively.

## Conclusions

This work illustrates that deep learning is required for efficient clustering in deformability cytometry, and provides a new technique for unsupervised clustering. Although simple dimensionality reduction techniques work well in a variety of areas, the work here shows that they are insufficient for the compression of complex data obtained from deformability cytometry. In addition, this work highlights the need for extensive VAE training for the purpose of feature learning. The potential of the SCAN loss in optimizing an encoder to classify unknown populations of cells is also explored using a novel unsupervised algorithm, DeepDeform. These improvements resulted in a model with moderate unsupervised clustering accuracy. Furthermore, when the predictions are thresholded, the accuracy of this model increases to 80%. A similar model could find applications in selecting rare, unknown groups of cells for further scrutiny, such as single cell RNA sequencing. Future work will entail the improvement of the interpretability of the DeepDeform. The features found most influential to DeepDeform can then be validated with downstream characterization, such as RNA sequencing or microscopy.

